# Toward Capturing the Exposome: Exposure Biomarker Variability and Co-Exposure Patterns in the Shared Environment

**DOI:** 10.1101/175513

**Authors:** Ming Kei Chung, Kurunthachalam Kannan, Germaine M. Buck Louis, Chirag J. Patel

**Affiliations:** Department of Biomedical Informatics Harvard Medical School Harvard University 10 Shattuck Street Boston, MA 02115; Division of Environmental Health Sciences Wadsworth Center, New York State Department of Health, and Department of Environmental Health Sciences The University at Albany Albany, New York 12201; Office of the Director Division of Intramural Population Health Research Eunice Kennedy Shriver National Institute for Child Health and Human Development The National Institutes of Health 6710b Rockledge Drive Bethesda, MD 20892 1-301-496-6155

**Keywords:** exposome, co-exposures, EDCs, combined exposure, POPs, metals, phthalates, persistent pollutants, mixtures, biomarkers, exposure assessment

## Abstract

**BACKGROUND:** Along with time, variation in the exposome is dependent on the location and sex of study participants. One specific factor that may influence exposure co-variations is a shared household environment.

**OBJECTIVES:** To examine the influence of shared household and partner’s sex in relation to the variation in 128 endocrine disrupting chemical (EDC) exposures among couples.

**METHODS:** In a cohort comprising 501 couples trying for pregnancy, we measured 128 (13 chemical classes) persistent and non-persistent EDCs and estimated 1) sex-specific differences; 2) variance explained by shared household; and 3) Spearman’s rank correlation coefficients (*r*_s_) for females, males, and couples’ exposures.

**RESULTS:** Sex was correlated with 8 EDCs including polyfluoroalkyl substances (PFASs) (*p* < 0.05). Shared household explained 43% and 41% of the total variance for PFASs and blood metals, respectively, but less than 20% for the remaining 11 EDC classes. Co-exposure patterns of the exposome were similar between females and males, with within-class *r*s higher for persistent and lower for non-persistent chemicals. Median *r*_s_s of polybrominated compounds and urine metalloids were 0.45 and 0.09, respectively, for females (0.41 and 0.08 for males), whereas lower *r*_s_s for these 2 classes were found for couples (0.21 and 0.04).

**CONCLUSIONS:** Overall, sex did not significantly affect EDC levels in couples. Individual, rather than shared environment, could be a major factor influencing the co-variation of 128 markers of the exposome. Correlations between exposures are lower in couples than in individual partners and have important analytical and sampling implications for epidemiological study.

## Introduction

Variation of environmental exposure levels in the population is a complex phenomenon and is influenced by factors shared by individuals — such as those within a household — and non-shared factors specific to individuals, such as their sex. One could apply the exposome paradigm (Wild 2005), a model to capture the totality of exposures from conception onwards, to comprehensively characterize the individual-level and shared differences of exposure. However, the exposome is a dynamic entity with variations across time (temporal) and place (spatial) (Wild et al. 2013) underscoring the importance of considering variability when assessing human health.

Instead of trying to “capture all” lifetime exposures, investigators can focus on critical and sensitive time windows in human development such as pregnancy, infancy, childhood and adolescence to reduce temporal complexity (Stingone et al. 2017). Furthermore, household-level ascertainments of exposure (i.e., sampling individuals from households) has been posited to be sufficient surrogates for all individuals in the household (Potera 2014). These assumptions may help characterize the exposome and study its time-dependent relationship with health outcomes. For example, the Human Early-Life Exposome (HELIX) project seeks to define the pregnancy and early-life exposomes and health (Vrijheid et al. 2014), while the EXPOsOMICS project has its conceptual framework of a life-course approach to a broader range of exposures (Vineis et al. 2017).

Another challenge includes the difficulty of interpreting exposure-disease associations due to the dense correlations among all exposures (Ioannidis 2016). The dense correlation pattern makes it hard to identify the directionality of the potential causality (Ioannidis et al. 2009). Second, correlations between exposures vary (e.g. absolute median correlation from almost 0 to above 0.5) and, thus, there is no universal scale to assess the biological significance (Patel and Ioannidis 2014). In addition, exposome-wide, or equivalently, environment-wide association studies (EWASs), assess all the associations between exposures and an outcome to identify potential etiologic signals (Manrai et al. 2017). The data-driven approach assumes little to no collinearity between environmental predictors, but it is almost impossible to select any single uncorrelated exposures out from the dense exposome. One strategy for addressing these analytical issues is to characterize the correlations in diverse cohorts to provide reference levels to gauge biological significance of associations.

We investigated whether cohabiting couples trying for pregnancy would have similar concentrations of endocrine disrupting chemicals (EDCs) given their shared households, and whether concentrations varied across partners in light of their individual exposures arising from other environments such as lifestyle, recreation or occupation. This avenue of study is important given that EDCs have been found to affect human fecundity and fertility (Hauser 2006; te Velde et al. 2010), though much of the available evidence relies on research conducted in either men or women but not couples. We utilized the Longitudinal Investigation of Fertility and the Environment (LIFE) Study to empirically assess couples’ shared and individual variations in a mixture of EDCs. We selected the LIFE Study because it has 13 classes of EDCs quantified in both partners of the couple in keeping with the couple based nature of human reproduction (Buck Louis et al. 2014, 2016; Patel et al. 2016). To meet our overarching aim, we characterize the dense correlation structure of couples’ EDC concentrations and then estimate shared and individual variability. Lastly, we discuss the implications of the findings in designing future exposome-related research.

## Methods

### Study Design and Cohort

Briefly, the LIFE Study enrolled 501 couples planning to discontinue contraception to become pregnant from 16 counties in Michigan and Texas, 2005–2009. Couples were followed daily until pregnant or up to 12 months of trying to become pregnant (infertile). Study participants were screened for eligibility based on a set of criteria and the complete details have been previously published (Buck Louis et al. 2011).

### Data Collection and Toxicologic Analysis

Following enrollment and completion of the baseline interview, couples provided preconception blood (20 mL) and urine (120 mL) samples for the quantification of both man-made and natural EDCs (e.g. phytoestrogens). We also included serum cotinine, a metabolite of nicotine, and have a total of 128 chemicals from 13 different classes (Table 1). Persistent EDCs included 4 classes of serum persistent organic compounds: 36 congeners of polychlorinated biphenyls (PCBs), 9 organochlorine pesticides (OCPs), 11 polybrominated chemicals [polybrominated diphenyl ethers (PBDEs) and 1 polybrominated biphenyl (PBB)], and 7 polyfluoroalkyl substances (PFASs), and were quantified by a single laboratory using published standard operating procedures (Kuklenyik et al. 2005; Sjödin et al. 2004).

**Table 1.** List of chemical classes and measured chemicals in the current study.

| EDC Class | No. | Individual Chemicals | Medium | LOQ |
| --- | --- | --- | --- | --- |
| Polychlorinated biphenyls (PCBs) | 36 | Congeners: 28, 44, 49, 52, 66, 74, 87, 99, 101, 105, 110, 114, 118, 128, 138, 146, 149, 151, 153, 156, 157, 167, 170, 172, 177, 178, 180, 183, 187, 189, 194, 195, 196, 201, 206, 209 | Serum | 1-3 pg/g, wet weight |
| Organochlorine pesticides (OCPs) | 9 | Hexachlorobenzene (HCB), β-hexachlorocyclohexane (β-HCH), γ-hexachlorocyclohexane (γ-HCH), oxychlorane, *trans*-nonachlor, *p,p'*-DDT, *o,p'*-DDT, *p,p'*-DDE, and mirex | Serum | 1-3 pg/g, wet weight |
| Polybrominated chemicals<sup>a</sup> | 11 | Brominated biphenyl (BB 153); brominated diphenyl ethers (BDEs) congeners: 17, 28, 47, 66, 85, 99, 100, 153, 154, 183 | Serum | 5 pg/g, wet weight |
| Polyfluoroalkyl substances (PFASs) | 7 | 2-(*N*-ethyl-perfluorooctane sulfonamido) acetate (Et-PFOSA-AcOH), 2-(*N*-methyl-perfluorooctane sulfonamido) acetate (Me-PFOSA-AcOH), perfluorodecanoate (PFDeA), perfluorononanoate (PFNA), perfluorooctane sulfonamide (PFOSA), perfluorooctane sulfonate (PFOS), and perfluorooctanoate (PFOA) | Serum | 0.04-0.1 ng/mL, wet weight |
| Phytoestrogens | 6 | Genistein, daidzein, *O*-desmethylangolensin, equol, enterodiol, and enterolactone | Urine | 0.2-0.6 ng/mL |
| Phthalate metabolites | 14 | Mono (3-carboxypropyl) phthalate (mCPP), monomethyl phthalate (mMP), monoethyl phthalate (mEP), mono (2-isobutyl phthalate) (miBP), mono-*n*-butyl phthalate (mBP), mono (2-ethyl-5-carboxyphenyl) phthalate (mECP), mono-[(2-carboxymethyl) hexyl] phthalate (mCMHP), mono (2-ethyl-5-oxohexyl) phthalate (mEOHP), mono (2-ethyl-5-hydroxyhexyl) phthalate (mEHHP), monocyclohexyl phthalate (mCHP), monobenzyl phthalate (mBzP), mono (2-ethylhexyl) phthalate (mEHP), mono-isononyl phthalate (mNP), and monooctyl phthalate (mOP) | Urine | 0.2-2 ng/mL |
| Phenols<sup>b</sup> | 6 | Total bisphenol A (BPA); benzophenones (BPs): 4-hydroxybenzophenone (4-OH-BP), 2,4-dihydroxybenzophenone (2,4-OH-BP), 2,2',4,4'-tetrahydroxybenzophenone (2,2',4,4'-OH-BP), 2-hydroxy-4-methoxybenzophenone (2-OH-4-MeO-BP), and 2,2'-dihydroxy-4-methoxybenzophenone (2,2'-OH-4-MeO-BP) | Urine | 0.02-0.05 ng/mL |
| Antimicrobial chemicals<sup>c</sup> | 12 | Triclosan (TCS) and triclocarban (TCC); parabens: methyl paraben (MP), ethyl paraben (EP), propyl paraben (PP), butyl paraben (BP), benzyl paraben (BzP), heptyl paraben (HP), 4-hydroxy benzoic acid (4-HB), 3,4-dihydroxy benzoic (3,4-DHB), methyl-protocatechuic acid (OH-Me-P), and ethyl-protocatechuic acid (OH-Et-P) | Urine | 0.02-0.05 ng/mL |
| Paracetamol & derivatives | 2 | Paracetamol and 4-aminophenol | Urine | 0.5 ng/mL and 0.25 ng/mL |
| Blood metals | 3 | Cadmium (Cd), lead (Pb), and mercury (Hg) | | 10 ng/L to 10 μg/L |
| Cotinine<sup>d</sup> | 1 | Cotinine | Serum | 0.01 ng/mL |
| Urine metals | 17 | Manganese (Mn), chromium (Cr), beryllium (Be), cobalt (Co), molybdenum (Mo), cadmium (Cd), tin (Sn), caesium (Cs), barium (Ba), nickel (Ni), copper (Cu), zinc (Zn), tungsten (W), platinum (Pt), thallium (Tl), lead (Pb), and uranium (U) | Urine | 10 ng/L to 10 μg/L |
| Urine metalloids | 4 | Selenium (Se), arsenic (As), antimony (Sb), and tellurium (Te) | Urine | 10 ng/L to 10 μg/L |

Note: LOQ, Limits of quantification.
<sup>a</sup>Polybrominated chemicals contain mostly PBDEs with one PBB. <sup>b</sup>Phenols contain mostly benzophenones with one BPA. <sup>c</sup>Antimicrobial chemicals contain mostly parabens with TCS and TCC. <sup>d</sup>Serum cotinine is not an EDC but included for comprehensive investigation.

Non-persistent ECDs included 5 classes of urinary non-persistent organic compounds: 6 phytoestrogens, 14 phthalate metabolites, 6 phenols [bisphenol A (BPA) and benzophenones (BPs)], 12 antimicrobial chemicals [parabens, triclosan (TCS) and triclocarban (TCC)], 2 paracetamol & derivatives, and were quantified by another laboratory using published standard operating procedures (Asimakopoulos et al. 2014; Guo et al. 2011; Mumford et al. 2015; Smarr et al. 2016). Other 3 classes of EDCs included 3 blood metals, 17 urinary metals, and 4 urinary metalloids (Bloom et al. 2015a).

Serum lipids were obtained by measuring total cholesterol, free cholesterol, triglycerides, and phospholipids by an enzymatic method (Akins et al. 1989) and we calculated the total serum lipids as described by Phillips et al. (1989). Creatinine was measured by a Roche/Hitachi Model 912 clinical analyzer (Dalla, TX, USA) using the Creatinine Plus Assay (Roche Diagnostics). Serum cotinine was quantified by isotope dilution tandem mass spectrometry (Bernert et al. 1997).

### Statistical Analysis

Our overall analytical scheme is shown in Figure 1. First, we adjusted each of the indicators of exposures for potential confounders in addition to total lipids (for lipophilic chemicals, *n* = 56) and creatinine (for urinary chemicals, *n* = 61) to reduce estimate variability and susceptibility to bias (Heavner et al. 2006; Schisterman et al. 2005). Specifically, we adjusted PCBs, OCPs, and polybrominated chemicals for total lipids and all other urinary chemicals for creatinine. Chemicals were log-transformed (x + 1) and continuous covariates were rescaled to have mean zero and unit variance (Figure 1A). After extracting the residuals, we calculated the Spearman’s rank correlation (*r*_s_) matrices for EDCs in females, males, and couples.

**Figure 1.**
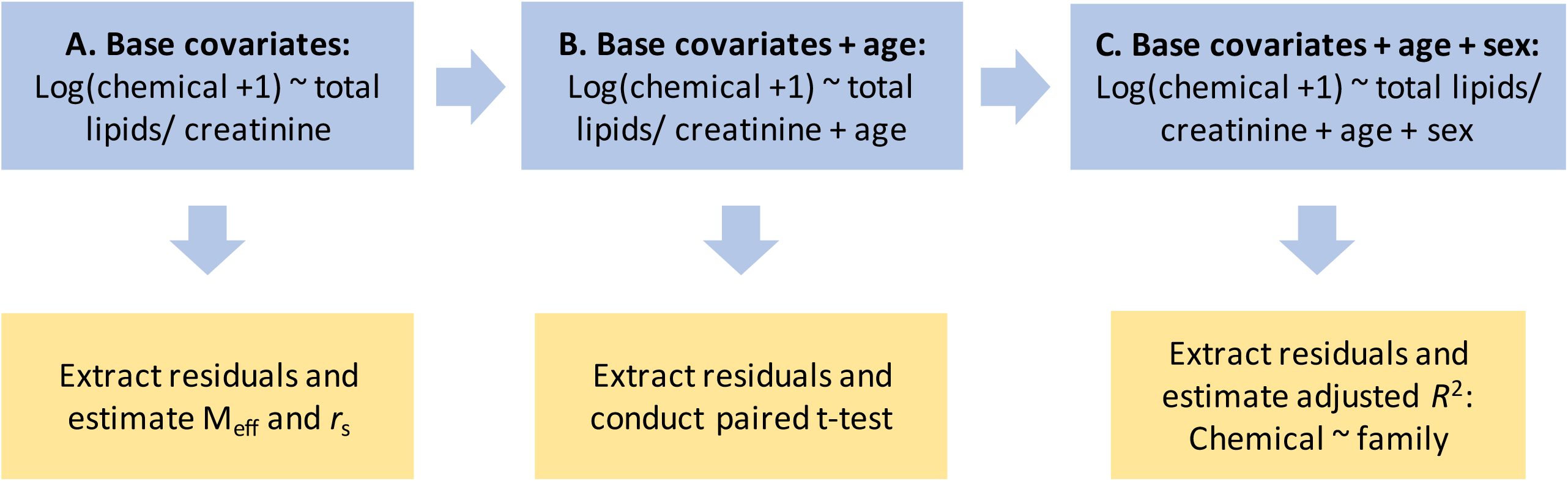
Analytical scheme to investigate the variability and correlations in this study. A) We first extract the residuals from a linear model after adjusting for the base covariates (total lipids or creatinine) to calculate the effective number of variables (M_eff_) and Spearman’s rank correlation (*r*_s_). B) Then, we used another linear model with an additional age variable to obtain residuals and conducted paired t-test to test the difference of biomarkers between females and males living in the same household. C) Afterward, we further adjusted for sex prior to extracting residuals to calculate the percentage of biomarker variance explained by the shared environment.

Adjusting procedures such as Bonferroni correction assume independence between independent variables. One way to resolve this analytical issue is to account for the correlation. To study the effect of correlations between exposures on the family-wise error rate, we calculated the effective number of variables (M_eff_) (Nyholt 2004) for estimating the Bonferroni adjusted *p* values with formula 1, where Var(λ_obs_) is the eigenvalue variance of the correlation matrix and M is the original number of variables.

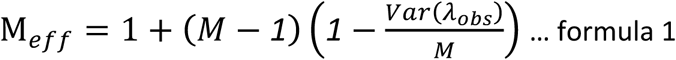

We estimated the sex-specific difference with a paired t-test (by household) after extracting the residuals from a linear model adjusted for age (Figure 1B). We used a similar approach to estimate the percentage of variance explained by the shared environment. However, sex and age variables were excluded in the adjustment step to isolate their effects (Figure 1C). Afterward, we extracted and regressed the residuals against the household variable to obtain the adjusted coefficient of determination (*R*^2^).

We computed concordance as the Pearson correlation coefficient (*r*) between the chemical relatedness *r*_s_ in this study and that in the 2003–2004 National Health and Nutrition Examination Survey (NHANES) (Patel and Ioannidis 2014) to assess generalizability of the co-exposure patterns. We estimated the concordance based on a total of 101 matched biomarkers between the 2 studies. We chose the 2003–2004 NHANES because 1) many of the chemicals measured in LIFE were also measured in NHANES and 2) by the same laboratory; and 3) the time period (2003–2004) is close to the beginning of recruitment for the LIFE Study.

We used all the instrument derived concentrations for the analyses (Richardson and Ciampi 2003; Schisterman et al. 2006). For missing values, we substituted them by multiple imputation, assuming a missing-at-random scenario (Louis et al. 2013). We conducted imputation based on the information from available demographic, previous history of clinical symptoms and all other chemical variables and created a total of 10 imputed data sets for males and females separately.

We visualized the correlations between exposures as exposome globe using the R package Circlize (v 0.3.1) (Gu et al. 2014). EDCs were sorted from lipophilic to hydrophilic to aid visual interpretation of the patterns. We combined the final estimates from imputations using Rubin’s method (Schafer 1999) and calculate the *p* values of correlations by permutation tests. To adjust for multiple testing, we used the false discovery rate (FDR) *q* values unless otherwise specified. We executed all analyses using the computing environment R (v 3.3.1) (R Core Team 2016). For reproducibility purpose, all analytic code is publicly available on GitHub via a MIT license (github.com/jakemkc/exposome_variability).

## Results

Important sociodemographic differences were observed between partners (Table 2). Overall, female partners were younger, had a lower body mass index (BMI) (< 25), consumed fewer alcoholic drinks, were less likely to report a hypertension or high cholesterol, and had lower serum cotinine and lipids and creatinine than male partners (*p* < 0.01).

**Table 2.** Sociodemographic and lifestyle characteristics of females and males in the LIFE Study.

| Characteristics | Females (*n* = 501) | Males (*n* = 501) |
| --- | --- | --- |
| Age (year) ** | 29.99 ± 4.14 | 31.77 ± 4.92 |
| BMI (kg/m<sup>2</sup>) ** |  |  |
| < 25 | 230 (46) | 84 (17) |
| ≥ 25 & < 30 | 136 (27) | 206 (41) |
| ≥ 30 | 135 (27) | 311 (62) |
| Non-Hispanic White | 396 (79) | 397 (79) |
| College graduate or higher * | 474 (95) | 457 (91) |
| Yearly income \$80,000 or over | 297 (59) | 298 (59) |
| Regular vigorous exercise in the past 12 months | 200 (40) | 211 (42) |
| Smoke at the time of study |  |  |
| No | 445 (89) | 440 (88) |
| Yes (No. of cigarettes on a typical day) |  |  |
| 1 - 3 | 19 (4) | 26 (5) |
| 4 - 6 | 8 (2) | 11 (2) |
| 7 - 10 | 15 (3) | 8 (2) |
| 11-15 | 7 (1) | 2 (0) |
| 16 - 25 | 3 (1) | 10 (2) |
| > 25 | 4 (1) | 4 (1) |
| ≥ 12 alcoholic drinks in the past 12 months ** | 374 (75) | 428 (85) |
| No. of alcoholic drinks on a typical occasion ** |  |  |
| 0 | 128 (26) | 73 (15) |
| 1 | 108 (22) | 63 (13) |
| 2 | 169 (34) | 150 (30) |
| 3 | 68 (14) | 99 (20) |
| 4 | 19 (4) | 62 (12) |
| 5 | 9 (2) | 54 (11) |
| History of diabetes | 6 (1) | 14 (3) |
| History of high blood pressure ** | 20 (4) | 52 (10) |
| History of high cholesterol ** | 41 (8) | 78 (16) |
| Serum cotinine (ng/mL)<sup>a</sup> ** | 0.62 ± 0.23 | 1.24 ± 2.17 |
| Serum total lipids (mg/dL)<sup>a</sup> ** | 2.00 ± 0.03 | 6.56 ± 0.26 |
| Urinary creatinine (mg/dL)<sup>a</sup> ** | 4.22 ± 0.86 | 4.76 ± 0.73 |

Note: BMI, body mass index. Values in mean ± SD or *n* (%).
\**p* < 0.05; \*\**p* < 0.01.
<sup>a</sup>log+1 transformed values.

Nine (7%) chemicals were correlated with partner’s sex (*p* < 0.05), i.e., cotinine, blood lead, mercury and cadmium; and 5 PFASs: perfluorodecanoate (PFDeA), perfluorononanoate (PFNA), perfluorooctane sulfonamide (PFOSA), perfluorooctane sulfonate (PFOS), and perfluorooctanoate (PFOA). Of note, findings were robust to the FDR with the exception of blood cadmium.

Figure 2 shows the boxplot summary of the variance explained by the shared environment. We estimated that two classes of chemicals, PFASs and blood metals, had higher levels of explained variance (median 0.43 and 0.41 respectively) than the others. For the rest of the 11 classes, median explained variances were ranged from 0.04 (phthalates) to 0.21 (cotinine). A few persistent organic compounds, namely PCB congener 28, PBDE congener 47, and *β*-hexachlorocyclohexane (*β*-HCH), had an explained variance over 50%.

**Figure 2.**
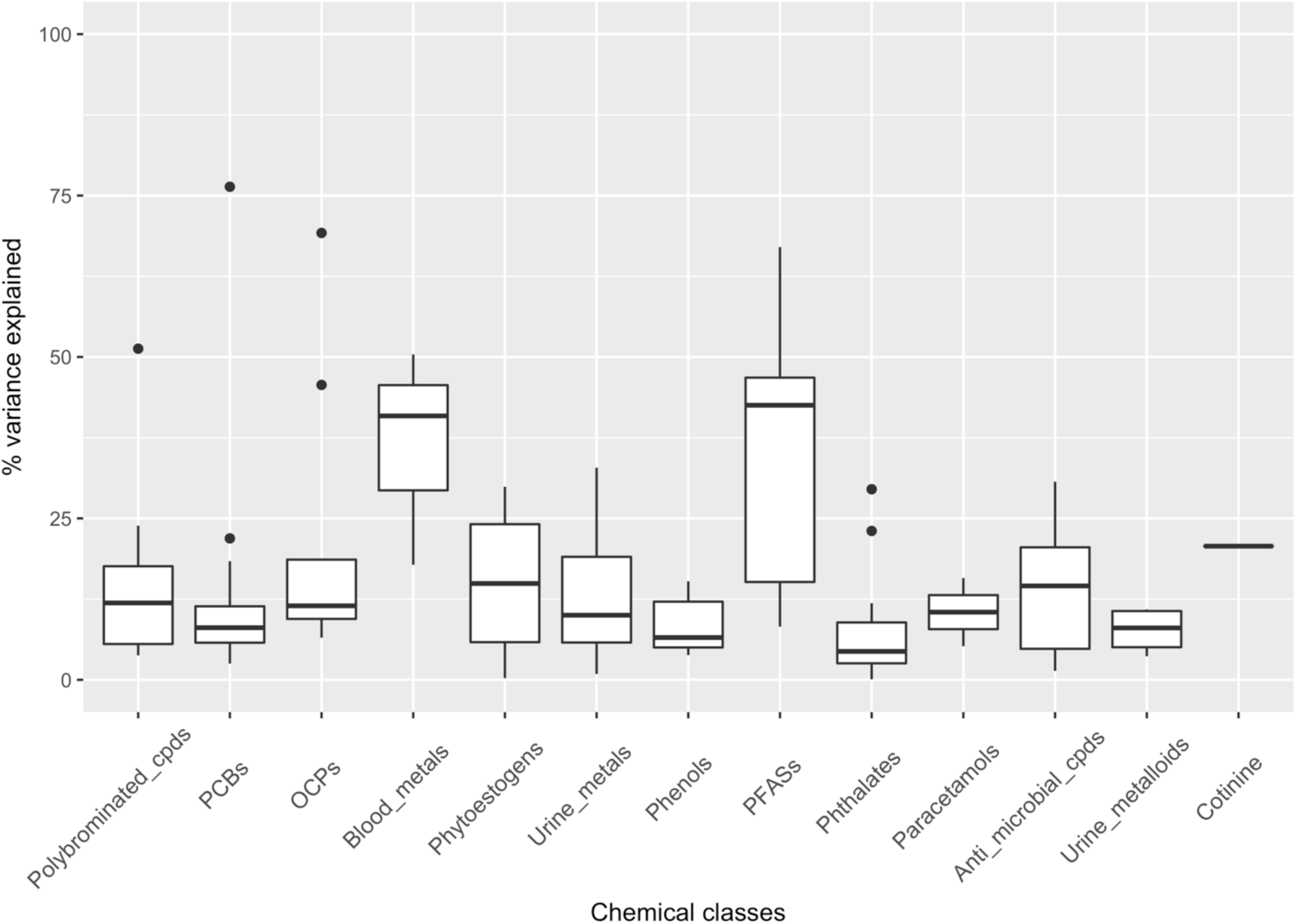
Summary of the percentage variance explained by the shared environment. Boxplots of the adjusted coefficient of determination (*R*^2^) within different chemical classes are shown. Interquartile range is not shown for Cotinine class because it contains only 1 compound. For each box, median and interquartile range are drawn and the whiskers are extended to the largest values within 1.5*interquartile range. Black dots denote correlations outside of the range covered by the whiskers.

The exposome globe (Patel and Manrai 2015) displays the *r*_s_s for females (right-half), for males (left-half), and for couples (across the left-right of the globe) (Figure 3). To assist interpretation, we only presented the *r*_s_s outside the range of −0.25 to 0.25 as lines connecting different parts on the track, and they represent less than 10% of all the *r*_s_s. For females, we observed two larger positively correlated “clusters” across EDC classes: A) a dense cluster with serum persistent organic compounds such as PCBs and OCPs (upper right of Figure 3); B) another loosely packed cluster with urinary EDCs such as phytoestrogens, phthalates, phenols, and antimicrobial compounds (lower right of Figure 3). Correlations between serum and urinary EDCs were mostly small and distributed between −0.25 and 0.25. For males, there were similar co-exposure patterns to females While we found similar correlations in the population of males and females separately, we found that correlations in couple living in the same household were, in fact, less densely packed and with values attenuated toward the null (Figure 3; see Figure S1).

**Figure 3.**
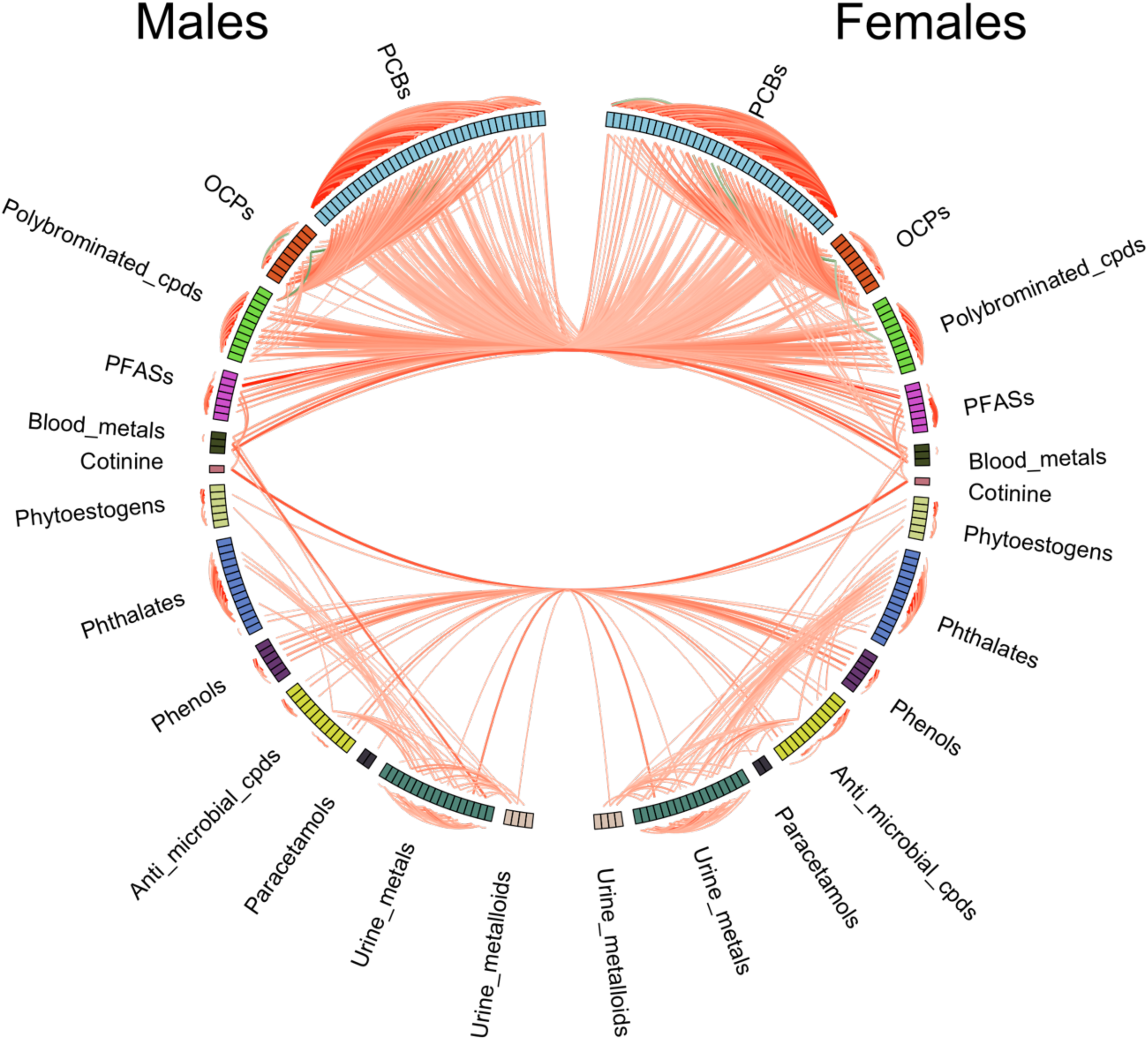
Exposome correlation globe showing the relationships between females, males and couples. Right-half represents biomarkers in females; left-half represents biomarkers in males. Only Spearman’s rank correlations greater than 0.25 and smaller than −0.25 were shown as connections in the globe. Red line denotes positive correlation and dark green line denotes negative one. Color intensity and line width are proportional to the size of the correlation. Within-class and between-class correlations are shown outside and inside of the track respectively. Correlations in couples are indicated by the lines linking females and males (i.e. crossing the vertical-half of the globe).

Summary of the within-class correlations as absolute magnitude is shown in Figure 4. For females (Figure 4A), we found that polybrominated compounds had the highest median correlation (*r*_s_ = 0.45) while urine metalloids had the lowest (*r*_s_ = 0.09). Across different chemical classes, persistent organic compounds such as polybrominated compounds, PCBs and OCPs had higher median correlations (0.45, 0.38, and 0.34 respectively). For the rest of the classes, the median correlations were all below 0.25. Males (Figure 4B) had similar within-class correlation distributions as were found in females. The class with highest and lowest median correlations were polybrominated compounds (*r*_s_ = 0.41) and paracetamols (*r*_s_ = 0.02) respectively.

**Figure 4.**
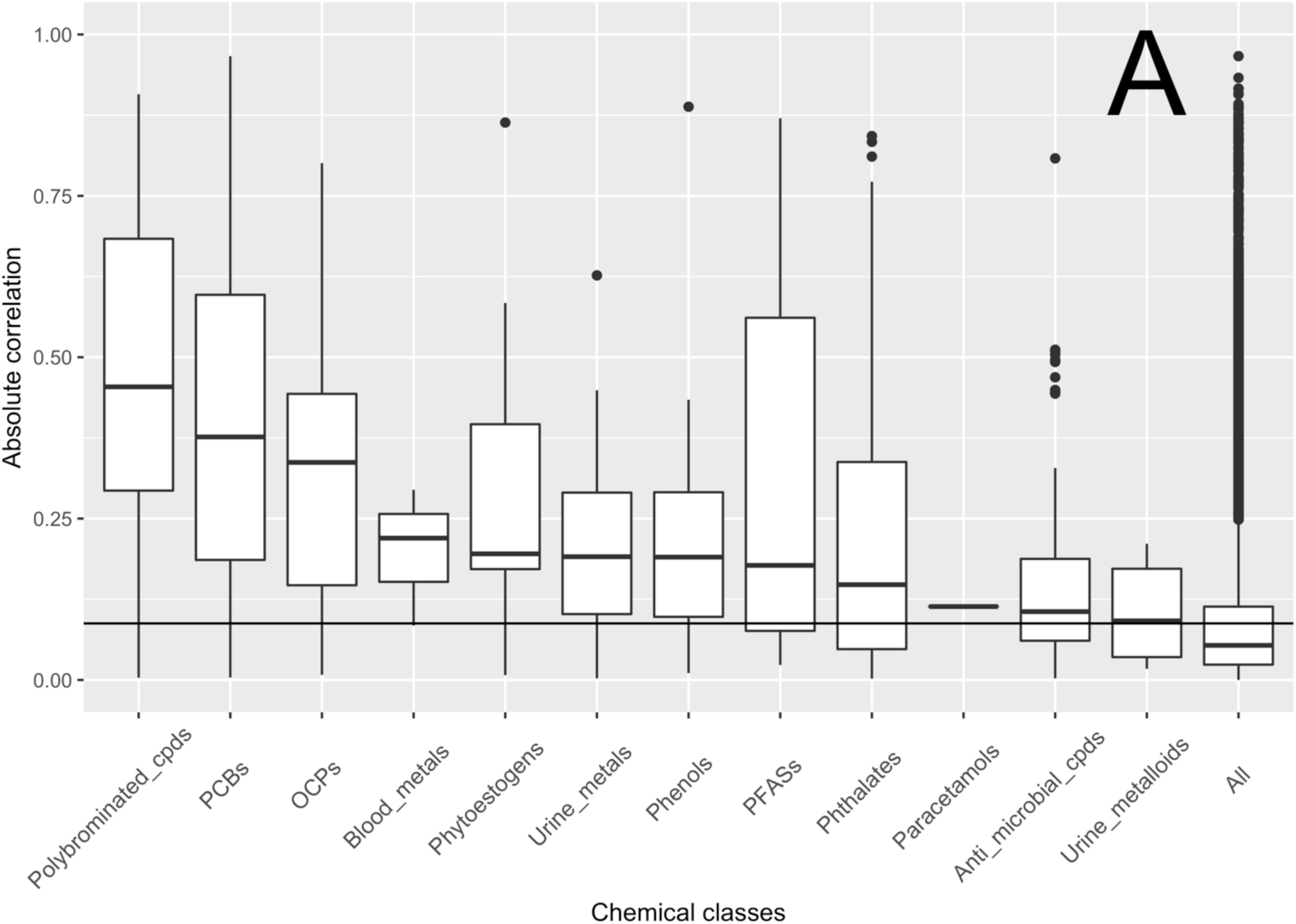
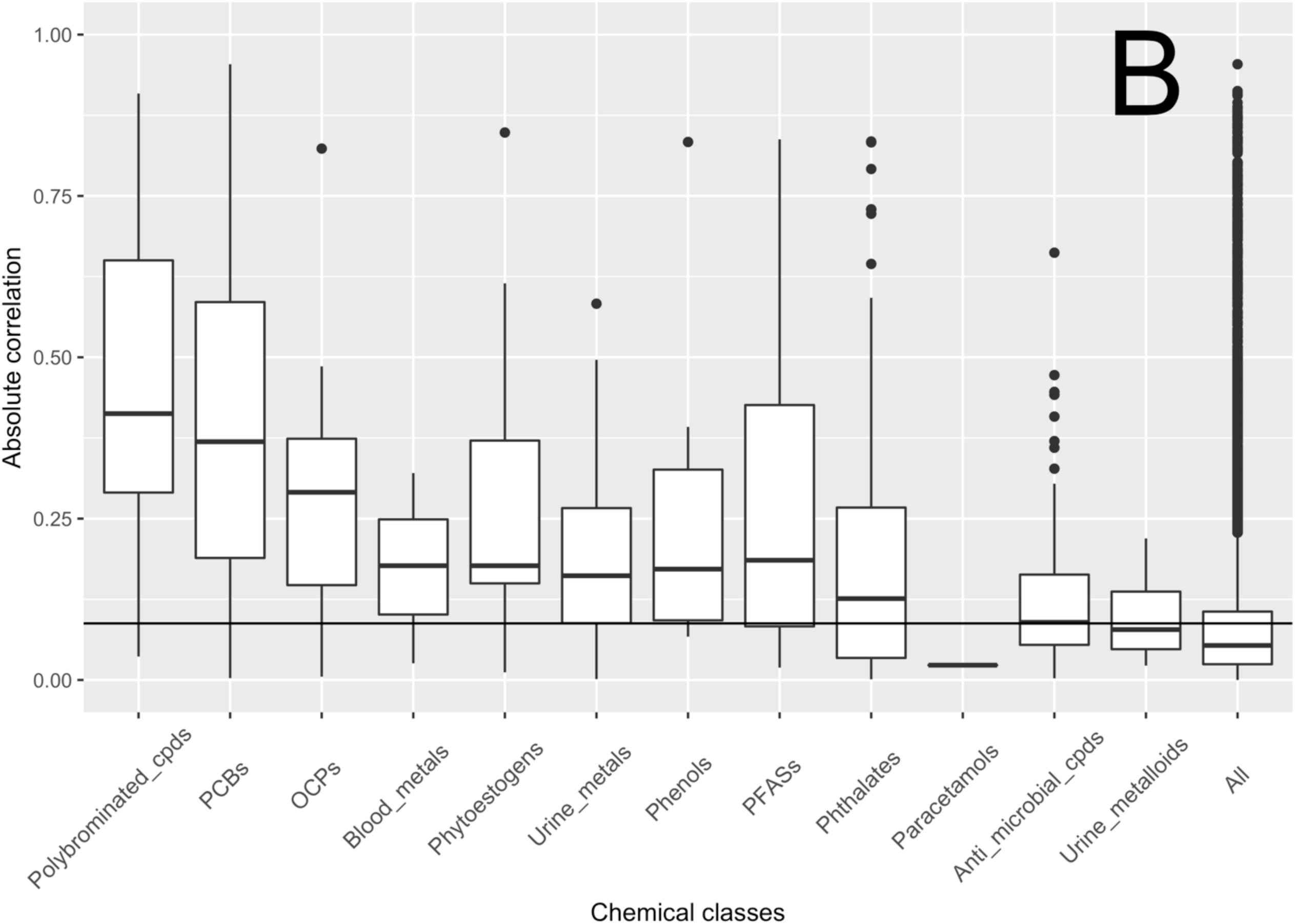
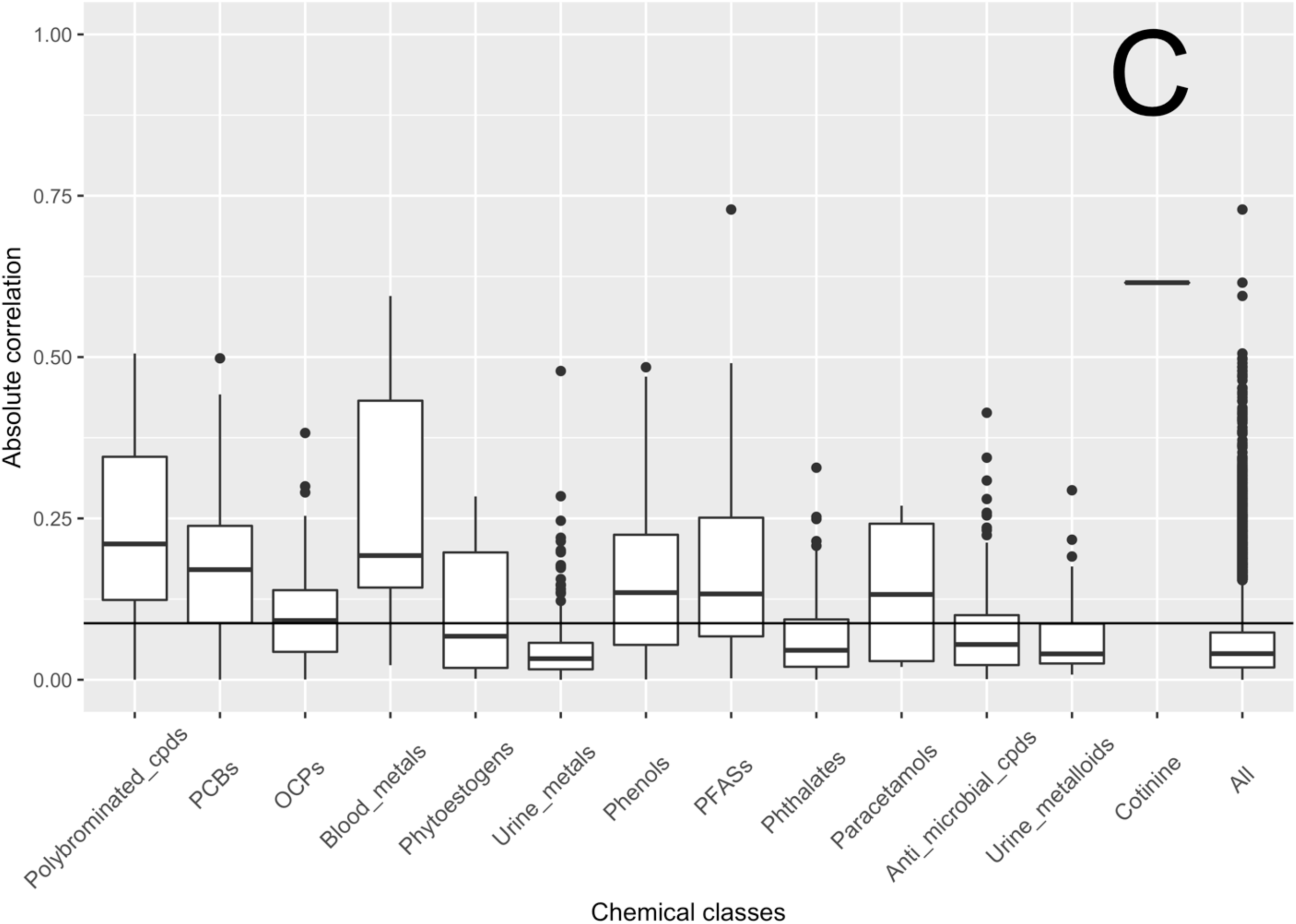
Boxplots of Spearman’s rank correlations (*r*_s_) within different chemical classes. A) Females; B) Males; and C) Couples. For couples, summary statistics were estimated with the full 128 x 128 correlation matrix instead of with the half triangle. Certain classes contain only 1 pair of correlation (paracetamols in females, paracetamols in males, and cotinine in couples). “All” represents the grouping by the correlation of all pairs of chemicals available. Horizontal line drawn across the chemical classes is equal to the 95^th^ percentile of the null distribution obtained from permuting the concentrations of all chemicals. Definition of whisker and black dot can be referred to the caption in Figure 2.

In contrast, we found a strong diminishing effects to the within-class correlations, both in terms of the median and interquartile range (IQR), in couples (Figure 4C). Comparing the couples with females and males, all chemical classes had the median correlations below 0.25 and urine metals were the class with greatest percentage drop (83%). We also observed a substantial reduction in the IQRs of the chemical classes. For example, the IQR of polybrominated compounds was 0.21, which corresponds to an over 35% drop relatively to females and males. We found that urine metals had the largest drop in IQR (over 77%). Looking more closely to the data, correlations between the same chemicals in couples (i.e. the diagonal of the correlation matrix of couples) were generally higher than that among the same class members. Because of the low number of exposure indicators in blood metals (3 chemicals), paracetamols (2 chemicals), and cotinine (1 chemical), the diminishing effect to within-class correlations was countered and thus the drop in medians and IQRs of these classes were not as prominent as others.

*P* values obtained from Bonferroni correction with M_eff_ were highly similar between females and males (Table 3). Both groups had similar M_eff_ values with a difference smaller than 1 variable, suggesting that the overall degree of correlations was also similar. After adjusting for the correlation, we found the total number of variables decreased from 128 to 112 for females and 113 for males.

**Table 3.** Number of effective variables for each chemical class after adjusting for the within-class correlation.

| Chemical class | Females |  |  |  | Males |  |  |
| --- | --- | --- | --- | --- | --- | --- | --- |
|  | M | M<sub>eff</sub> | M<sub>diff</sub> | P value | M<sub>eff</sub> | M<sub>diff</sub> | P value |
| Serum persistent organic compounds |  |  |  |  |  |  |  |
| Polychlorinated biphenyls (PCBs) | 36 | 28.38 | 7.62 | 0.002 | 28.69 | 7.31 | 0.002 |
| Organochlorine pesticides (OCPs) | 9 | 7.94 | 1.06 | 0.006 | 8.17 | 0.83 | 0.006 |
| Polybrominated chemicals | 11 | 8.22 | 2.78 | 0.006 | 8.38 | 2.62 | 0.006 |
| Polyfluoroalkyl substances (PFASs) | 7 | 6.13 | 0.87 | 0.008 | 6.25 | 0.75 | 0.008 |
| Urinary non-persistent organic compounds |  |  |  |  |  |  |  |
| Phytoestrogens | 6 | 5.34 | 0.66 | 0.009 | 5.38 | 0.62 | 0.009 |
| Phthalate metabolites | 14 | 12.72 | 1.28 | 0.004 | 12.87 | 1.13 | 0.004 |
| Phenols | 6 | 5.48 | 0.52 | 0.009 | 5.51 | 0.49 | 0.009 |
| Antimicrobial chemicals | 12 | 11.38 | 0.62 | 0.004 | 11.60 | 0.40 | 0.004 |
| Paracetamol & derivatives | 2 | 1.99 | 0.01 | 0.025 | 2.00 | 0.00 | 0.025 |
| Others |  |  |  |  |  |  |  |
| Blood metals | 3 | 2.91 | 0.09 | 0.017 | 2.91 | 0.09 | 0.017 |
| Serum cotinine | 1 | 1.00 | 0.00 | 0.050 | 1.00 | 0.00 | 0.050 |
| Urine metals | 17 | 16.13 | 0.87 | 0.003 | 16.22 | 0.78 | 0.003 |
| Urine metalloids | 4 | 3.95 | 0.05 | 0.013 | 3.95 | 0.05 | 0.013 |
| Total | 128 | 111.57 | 16.43 | 0.0004 | 112.94 | 15.06 | 0.0004 |

Note: M, number of variables in the corresponding chemical classes; M<sub>eff</sub>, number of effective variables after taking account of within-class correlation; M<sub>diff</sub>, the different between M and M<sub>eff</sub>; P value, Bonferroni adjusted p values as 0.05/M<sub>eff</sub>.

We estimated the concordances between correlations in the LIFE study (as females, males and couples) and the 2003-2004 NHANES in Table 4. Overall, correlations *r* varied greatly between chemical groups, from −0.78 to 0.98. Certain chemical groups had small sample size (*n* ≤ 6) and could cause low and inverse correlations. However, correlations *r* were more consistent and comparable when we discarded group information and considered all chemicals as a whole (females: 0.88; males: 0.84; couples: 0.67); therefore, we conclude the exposure correlation patterns captured in LIFE are comparable to that in the U.S. population.

**Table 4.** Correlations of chemical relatedness between 2003–2004 NHANES and current study by different chemical classes.

| Chemical class<sup>a</sup> | Females |  | Males |  | Couples |  |
| --- | --- | --- | --- | --- | --- | --- |
|  | Pearson *r* | *n* | Pearson *r* | *n* | Pearson *r* | *n* |
| Blood_metals | 0.07 | 3 | 0.13 | 3 | -0.04 | 6 |
| OCPs | 0.69 | 36 | 0.51 | 36 | 0.25 | 72 |
| PCBs | 0.88 | 561 | 0.88 | 561 | 0.77 | 1122 |
| PFASs | 0.31 | 21 | 0.29 | 21 | 0.32 | 42 |
| Phthalates | 0.90 | 78 | 0.89 | 78 | 0.34 | 156 |
| Phytoestrogens | 0.98 | 15 | 0.97 | 15 | 0.86 | 30 |
| Polybrominated_cpds | 0.76 | 55 | 0.74 | 55 | 0.77 | 110 |
| Urine_metalloids | 0.41 | 3 | -0.78 | 3 | 0.34 | 6 |
| Urine_metals | 0.82 | 55 | 0.78 | 55 | -0.01 | 110 |
| Total | 0.84 | 827 | 0.84 | 827 | 0.67 | 1654 |

Note: Pearson *r*, Pearson correlation coefficients of the chemical relatedness (spearman correlations of chemicals) between National Health and Nutrition Examination Survey (NHANES) and this study; *n*, sample size.
<sup>a</sup>Only 9 classes are shown because not all chemicals in 13 chemical classes in this study could be matched with NHANES.

## Discussion

### Findings

Understanding the co-exposure patterns is an important step toward investigating the joint health effects of chemical mixtures and for statistical design of exposome-related investigations. We describe our high level findings here. First, exposure levels of one individual in the household were not correlated with another individual in the same household. Second, the percentages of significant *r*_s_ (*q* < 0.05) in males and females were 25.3 and 23.1 respectively, in contrast to only 9.5% between couples in the same household (Figure 4). Chemical correlations in a household setting were concordant to those in 03–04 NHANES, indicating reproducible co-exposure correlations with respect to the patterns sought in a generalized and non-institutionalized U.S. population.

Although couples in our study potentially shared a large degree of dietary and indoor environmental factors, their exposures were only modestly correlated (low *r*_s_). We believe that there are two additional factors affecting the familial co-exposure patterns in our investigation. The first one is concerned with how long the couples have been living together. In the U.S., the median age of first marriage is over 24 since 2000 (U.S. Census Bureau 2016), and newly married couples could show a greater discordance in chemical co-exposure relationships due to a different pre-marriage exposures. The second factor is potential physiological dampening of exposure variability related to the half-life of the target chemicals (Makey et al. 2014; Rappaport and Kupper 2008). Polybrominated compounds, PCBs, and OCPs are persistent chemicals with high lipophilicity and longer half-lives (on the order of years). Their serum concentrations are integrated over a period of time and not completely associated with recent exposure (Aylward et al. 2014). Since most of the couples recruited in LIFE Study were living together, we claim this phenomenon can explain the drop in *r*_s_ of persistent chemicals relative to the short-lived urinary chemicals (such as phthalate metabolites, whose half-lives are on the order of days) in couples (Figure 4).

Wu et al. (2015) conducted a household-based study and measured serum PBDE levels in children and parents and reported the *r*_s_ between child and parent were in the range of 0.66 to 0.74 (median = 0.68, *n* = 68) for a number of PBDE compounds. The pairs shared a substantial portion of genes, diet, and living environment and they found that for the latter 2 components, measured as floor-wipe PBDEs, canned meat, tuna and whitefish, were predictive of serum PBDEs. Furthermore, they found higher *r*_s_ of PBDEs between older couples (age ≥ 55, range = 0.45 to 0.78, median = 0.72). These findings are consistent with our claim.

Persistent organic chemicals are known to cause adverse health effects and are prioritized by the U.S. Centers for Disease Control and Prevention for health monitoring (Li et al. 2006). Many of them had wide applications in electrical/electronic equipment, agricultural chemicals, and furniture (Dodson et al. 2012; Whitehead et al. 2011). Although other emerging EDCs such as BPA and phthalates have short half-lives, they have extensive modern applications in cosmetics and consumer products (Bloom et al. 2015b; Buck Louis et al. 2014; Louis et al. 2014; Smarr et al. 2017). The ubiquity of these chemicals in different microenvironments such as schools and offices suggests that household environment alone is not a major contributor to body burden, consistent with our results (Figure 2). PFASs and blood metals had higher variance explained by the shared environment than other chemical classes and we believe that this could be related to the lifestyle of the subjects in the LIFE cohort (e.g. PFAS exposures via food items and personal care products).

### Strength and Limitations

Our investigation includes 128 chemical biomarkers with diverse physicochemical properties that span from persistent lipophilic to non-persistent hydrophilic chemicals and is one of the first attempts to systematically characterize their correlations in a household setting. However, we do not know how long the couples have been living together at the baseline of the study to quantitatively assess how this factor affects co-exposure patterns. Also, we only collected biological samples at the baseline; thus, it is not possible to study how exposure levels and co-exposure patterns change longitudinally with time, which could be an important piece of information for assessing fecundity outcome and chemical exposures.

### Analytical and Sampling Implications for Exposome-Wide Investigations

Our findings have implications for high-throughput association tests between correlated exposures and health outcomes and phenotypes. One of the typical approaches adjusting for multiplicity in EWASs is controlling the family-wise error rate (e.g. Bonferroni correction). However, the tests are assumed to be independent. Nyholt (2004) provided one solution to address correlation in Bonferroni correction by calculating the effective number of variables (M_eff_, formula 1), which relies on identifying the number of variables, M, and estimating the eigenvalue variance, Var(λ_obs_) of the correlation matrix. For example, conducting an EWAS of a response with respect to the chemical class PCBs containing 36 congeners, one can calculate the correlation matrix (M = 36) and estimate the associated Var(λ_obs_) to obtain M_eff_ (Table 3). A new significance level for this set of comparisons will be α/Meff, which is less stringent than Bonferroni correction because M_eff_ wil be smaller than M if correlations exist between congeners. Benjamini and Yekutieli (2001) and Fan et al. (2012) documented a procedure that considers the correlation structure for better controlling the FDR.

In the future, exposome-related investigations, will include many more variables than ever before with the emergence of highly sensitive high-resolution mass spectrometry techniques (Jones 2016) and availability of large data sets from reproducibility initiatives (Manrai et al. 2017). Attempts to assess health outcome associated with a number of exposures may increase *R*^2^ and lead to “overfitting” (Hawkins 2004). Additionally, dense correlations among exposures indicate that multicollinearity may also influence the reliability of the association size in multiple regression or even potential confounding. We claim that these analytic challenges could be ameliorated through understanding co-exposure patterns. For example, statistical approaches for evaluating the effects of mixture exposures such as principal component analysis generally involve dimensionality reduction that relies on estimating the correlation structure to reduce the number of exposures being considered prior to analysis but are difficult to interpret (Taylor et al. 2016). Other regression approaches, such as the Elastic Net (Zou and Hastie 2005) can consider highly co-linear exposure variables while giving coefficients that are similar to that delivered from a typical regression model; however, inferential estimates (e.g. *p* values and confidence intervals) are still in development (Taylor and Tibshirani 2015).

Capturing population-level exposure variability — and the demographic variables that are associated with the variation, such as sex, location, and time — is a grand ambition in the exposome concept (Manrai et al. 2017). Given people spend over 90% of their time indoor and more than 12 hours a day at home (bls.gov/tus), household samples (e.g. such as house dust) might be a reasonable surrogate to represent home exposures to family members who share the same living environment. While sampling the household is a tempting approach because of the simplicity and cost-savings over personal measurement, it may fail to catch a significant fraction of exposure variability in the population as we found that shared environment could explain a small percentage of biomarker variance (Figure 2). For epidemiological investigation that sample study participants who are nested in a unit, for example, schools or homes, conducting preliminary measurements to assess the influence of shared environment is one of the ways to justify unit-based measurement.

Finally, correlations of chemical mixtures can also be an important tool used in exposure science and cumulative risk assessment as groups of correlated chemicals are often released from a single source (e.g. power plant and vehicular exhaust (Ravindra et al. 2008). Thus, studying the co-exposure patterns is the first step to identify the sources prior to more in-depth source apportionment methods such as positive matrix factorization and chemical mass balance receptor models (Rizzo and Scheff 2007). In cumulative risk assessment, dose addition is one of the common approaches to estimate the risks from mixture exposures by assuming a shared toxicity mechanism between chemicals (e.g. binding to the same receptor) (Chen et al. 2001). Knowing the co-exposure correlations could be a first step toward identification of exposures with similar physicochemical properties to guide follow-up investigations.

## Conclusions

While we observed similar co-exposure patterns between females and males, the correlations were much lower in couples. Our analyses empirically demonstrate that shared environment explains less than 20% of the biomarker variance in 11 out of 13 EDC classes. The influence of shared environment to EDC levels is likely conditional on 1) the duration of residence of the subjects and 2) the lipophilicity and persistency of the chemicals. These factors should be considered when using surrogate measurement to assess the exposures of family members.

## Acknowledgement

The authors declare they have no actual or potential competing financial interests. This study is supported by the NIH grants (ES025052 and ES023504), and the Intramural Research Program of the *Eunice Kennedy Shriver* National Institute of Child Health and Human Development (contracts # N01-HD-3-3355; N01-HD-3-3356; N01-HD-3-3358; HHSN27500002; HHSN27500003; HHSN27500006)

## References

Akins JR, Waldrep K, Bernert JT. 1989. The estimation of total serum lipids by a completely enzymatic “summation” method. Clin. Chim. Acta 184: 219–226.

Asimakopoulos AG, Wang L, Thomaidis NS, Kannan K. 2014. A multi-class bioanalytical methodology for the determination of bisphenol A diglycidyl ethers, p-hydroxybenzoic acid esters, benzophenone-type ultraviolet filters, triclosan, and triclocarban in human urine by liquid chromatography-tandem mass spectrometry. J. Chromatogr. A 1324: 141–148.

Aylward LL, Hays SM, Smolders R, Koch HM, Cocker J, Jones K, et al. 2014. Sources of variability in biomarker concentrations. J. Toxicol. Environ. Health B Crit. Rev. 17: 45–61.

Benjamini Y, Yekutieli D. 2001. The Control of the False Discovery Rate in Multiple Testing under Dependency. Ann. Stat. 29: 1165–1188.

Bernert JT Jr, Turner WE, Pirkle JL, Sosnoff CS, Akins JR, Waldrep MK, et al. 1997. Development and validation of sensitive method for determination of serum cotinine in smokers and nonsmokers by liquid chromatography/atmospheric pressure ionization tandem mass spectrometry. Clin. Chem. 43: 2281–2291.

Bloom MS, Buck Louis GM, Sundaram R, Maisog JM, Steuerwald AJ, Parsons PJ. 2015a. Birth outcomes and background exposures to select elements, the Longitudinal Investigation of Fertility and the Environment (LIFE). Environ. Res. 138: 118–129.

Bloom MS, Whitcomb BW, Chen Z, Ye A, Kannan K, Buck Louis GM. 2015b. Associations between urinary phthalate concentrations and semen quality parameters in a general population. Hum. Reprod. 30: 2645–2657.

Buck Louis GM, Barr DB, Kannan K, Chen Z, Kim S, Sundaram R. 2016. Paternal exposures to environmental chemicals and time-to-pregnancy: overview of results from the LIFE study. Andrology 4: 639–647.

Buck Louis GM, Schisterman EF, Sweeney AM, Wilcosky TC, Gore-Langton RE, Lynch CD, et al. 2011. Designing prospective cohort studies for assessing reproductive and developmental toxicity during sensitive windows of human reproduction and development – the LIFE Study. Paediatr. Perinat. Epidemiol. 25: 413–424.

Buck Louis GM, Sundaram R, Sweeney AM, Schisterman EF, Maisog J, Kannan K. 2014. Urinary bisphenol A, phthalates, and couple fecundity: the Longitudinal Investigation of Fertility and the Environment (LIFE) Study. Fertil. Steril. 101: 1359–1366.

Chen JJ, Chen YJ, Rice G, Teuschler LK, Hamernik K, Protzel A, et al. 2001. Using dose addition to estimate cumulative risks from exposures to multiple chemicals. Regul. Toxicol. Pharmacol. 34: 35–41.

Dodson RE, Perovich LJ, Covaci A, Van den Eede N, Ionas AC, Dirtu AC, et al. 2012. After the PBDE phase-out: a broad suite of flame retardants in repeat house dust samples from California. Environ. Sci. Technol. 46: 13056–13066.

Fan J, Han X, Gu W. 2012. Estimating False Discovery Proportion Under Arbitrary Covariance Dependence. J. Am. Stat. Assoc. 107: 1019–1035.

Guo Y, Alomirah H, Cho H-S, Minh TB, Mohd MA, Nakata H, et al. 2011. Occurrence of phthalate metabolites in human urine from several Asian countries. Environ. Sci. Technol. 45: 3138–3144.

Gu Z, Gu L, Eils R, Schlesner M, Brors B. 2014. circlize Implements and enhances circular visualization in R. Bioinformatics 30: 2811–2812.

Hauser R. 2006. The environment and male fertility: recent research on emerging chemicals and semen quality. Semin. Reprod. Med. 24: 156–167.

Hawkins DM. 2004. The problem of overfitting. J. Chem. Inf. Comput. Sci. 44: 1–12.

Heavner DL, Morgan WT, Sears SB, Richardson JD, Byrd GD, Ogden MW. 2006. Effect of creatinine and specific gravity normalization techniques on xenobiotic biomarkers in smokers' spot and 24-h urines. J. Pharm. Biomed. Anal. 40: 928–942.

Ioannidis JPA. 2016. Exposure-wide epidemiology: revisiting Bradford Hill. Stat. Med. 35: 1749–1762.

Ioannidis JPA, Loy EY, Poulton R, Chia KS. 2009. Researching genetic versus nongenetic determinants of disease: a comparison and proposed unification. Sci. Transl. Med. 1: 7ps8.

Jones DP. 2016. Sequencing the exposome: A call to action. Toxicol Rep 3: 29–45.

Kuklenyik Z, Needham LL, Calafat AM. 2005. Measurement of 18 perfluorinated organic acids and amides in human serum using on-line solid-phase extraction. Anal. Chem. 77: 6085–6091.

Li QQ, Loganath A, Chong YS, Tan J, Obbard JP. 2006. Persistent organic pollutants and adverse health effects in humans. J. Toxicol. Environ. Health A 69: 1987–2005.

Louis GMB, Kannan K, Sapra KJ, Maisog J, Sundaram R. 2014. Urinary concentrations of benzophenone-type ultraviolet radiation filters and couples' fecundity. Am. J. Epidemiol. 180: 1168–1175.

Louis GMB, Sundaram R, Schisterman EF, Sweeney AM, Lynch CD, Gore-Langton RE, et al. 2013. Persistent environmental pollutants and couple fecundity: the LIFE study. Environ. Health Perspect. 121: 231–236.

Makey CM, McClean MD, Sjödin A, Weinberg J, Carignan CC, Webster TF. 2014. Temporal variability of polybrominated diphenyl ether (PBDE) serum concentrations over one year. Environ. Sci. Technol. 48: 14642–14649.

Manrai AK, Cui Y, Bushel PR, Hall M, Karakitsios S, Mattingly CJ, et al. 2017. Informatics and Data Analytics to Support Exposome-Based Discovery for Public Health. Annu. Rev. Public Health 38: 279–294.

Mumford SL, Kim S, Chen Z, Boyd Barr D, Buck Louis GM. 2015. Urinary Phytoestrogens Are Associated with Subtle Indicators of Semen Quality among Male Partners of Couples Desiring Pregnancy. J. Nutr. 145: 2535–2541.

Nyholt DR. 2004. A simple correction for multiple testing for single-nucleotide polymorphisms in linkage disequilibrium with each other. Am. J. Hum. Genet. 74: 765–769.

Patel CJ, Ioannidis JPA. 2014. Placing epidemiological results in the context of multiplicity and typical correlations of exposures. J. Epidemiol. Community Health 68: 1096–1100.

Patel CJ, Manrai AK. 2015. Development of exposome correlation globes to map out environment-wide associations. Pac. Symp. Biocomput. 231–242.

Patel CJ, Sundaram R, Buck Louis GM. 2016. A data-driven search for semen-related phenotypes in conception delay. Andrology 5: 95–102.

Phillips DL, Pirkle JL, Burse VW, Bernert JT Jr, Henderson LO, Needham LL. 1989. Chlorinated hydrocarbon levels in human serum: effects of fasting and feeding. Arch. Environ. Contam. Toxicol. 18: 495–500.

Potera C. 2014. The HELIX Project: tracking the exposome in real time. Environ. Health Perspect. 122: A169.

Rappaport SM, Kupper LL. 2008. Quantitative exposure assessment. El Cerrito, CA:S. Rappaport.

Ravindra K, Sokhi R, Van Grieken R. 2008. Atmospheric polycyclic aromatic hydrocarbons: Source attribution, emission factors and regulation. Atmos. Environ. 42: 2895–2921.

R Core Team. 2016. R: A Language and Environment for Statistical Computing.

Richardson DB, Ciampi A. 2003. Effects of exposure measurement error when an exposure variable is constrained by a lower limit. Am. J. Epidemiol. 157: 355–363.

Rizzo MJ, Scheff PA. 2007. Utilizing the Chemical Mass Balance and Positive Matrix Factorization models to determine influential species and examine possible rotations in receptor modeling results. Atmos. Environ. 41: 6986–6998.

Schafer JL. 1999. Multiple imputation: a primer. Stat. Methods Med. Res. 8: 3–15.

Schisterman EF, Vexler A, Whitcomb BW, Liu A. 2006. The limitations due to exposure detection limits for regression models. Am. J. Epidemiol. 163: 374–383.

Schisterman EF, Whitcomb BW, Louis GMB, Louis TA. 2005. Lipid adjustment in the analysis of environmental contaminants and human health risks. Environ. Health Perspect. 113: 853–857.

Sjödin A, Jones RS, Lapeza CR, Focant J-F, McGahee EE 3rd, Patterson DG Jr. 2004. Semiautomated high-throughput extraction and cleanup method for the measurement of polybrominated diphenyl ethers, polybrominated biphenyls, and polychlorinated biphenyls in human serum. Anal. Chem. 76: 1921–1927.

Smarr MM, Grantz KL, Sundaram R, Maisog JM, Honda M, Kannan K, et al. 2016. Urinary paracetamol and time-to-pregnancy. Hum. Reprod. 31: 2119–2127.

Smarr MM, Sundaram R, Honda M, Kannan K, Louis GMB. 2017. Urinary Concentrations of Parabens and Other Antimicrobial Chemicals and Their Association with Couples' Fecundity. Environ. Health Perspect. 125: 730–736.

Stingone JA, Buck Louis GM, Nakayama SF, Vermeulen RCH, Kwok RK, Cui Y, et al. 2017. Toward Greater Implementation of the Exposome Research Paradigm within Environmental Epidemiology. Annu. Rev. Public Health 38: 315–327.

Taylor J, Tibshirani RJ. 2015. Statistical learning and selective inference. Proc. Natl. Acad. Sci. U. S. A. 112: 7629–7634.

Taylor KW, Joubert BR, Braun JM, Dilworth C, Gennings C, Hauser R, et al. 2016. Statistical Approaches for Assessing Health Effects of Environmental Chemical Mixtures in Epidemiology: Lessons from an Innovative Workshop. Environ. Health Perspect. 124: A227–A229.

Velde E, Burdorf A, Nieschlag E, Eijkemans R, Kremer JAM, Roeleveld N, et al. 2010. Is human fecundity declining in Western countries? Hum. Reprod. 25: 1348–1353.

U.S. Census Bureau. 2016. Current Population Survey: Annual Social and Economic Supplement.

Vineis P, Chadeau-Hyam M, Gmuender H, Gulliver J, Herceg Z, Kleinjans J, et al. 2017. The exposome in practice: Design of the EXPOsOMICS project. Int. J. Hyg. Environ. Health 220: 142–151.

Vrijheid M, Slama R, Robinson O, Chatzi L, Coen M, van den Hazel P, et al. 2014. The human early-life exposome (HELIX): project rationale and design. Environ. Health Perspect. 122: 535–544.

Whitehead T, Metayer C, Buffler P, Rappaport SM. 2011. Estimating exposures to indoor contaminants using residential dust. J. Expo. Sci. Environ. Epidemiol. 21: 549–564.

Wild CP. 2005. Complementing the Genome with an “Exposome”: The Outstanding Challenge of Environmental Exposure Measurement in Molecular Epidemiology. Cancer Epidemiol. Biomarkers Prev. 14: 1847–1850.

Wild CP, Scalbert A, Herceg Z. 2013. Measuring the exposome: a powerful basis for evaluating environmental exposures and cancer risk. Environ. Mol. Mutagen. 54: 480–499.

Wu XM, Bennett DH, Moran RE, Sjödin A, Jones RS, Tancredi DJ, et al. 2015. Polybrominated diphenyl ether serum concentrations in a Californian population of children, their parents, and older adults: an exposure assessment study. Environ. Health 14: 23.

Zou H, Hastie T. 2005. Regularization and variable selection via the elastic net. J. R. Stat. Soc. Series B Stat. Methodol. 67: 301–320.

